# Single-nucleus RNA sequencing shows convergent evidence from different cell types for altered synaptic plasticity in major depressive disorder

**DOI:** 10.1101/384479

**Authors:** Corina Nagy, Malosree Maitra, Arnaud Tanti, Matthew Suderman, Jean-Francois Théroux, Naguib Mechawar, Jiannis Ragoussis, Gustavo Turecki

## Abstract

Major depressive disorder (MDD) is a complex illness that involves the interaction of different brain systems, pathways, and cell types. Past molecular studies of MDD relied on cellular homogenates of post-mortem brain tissue, making it impossible to determine gene expression changes within individual cells. Using single-cell transcriptomics, we examined almost 80,000 nuclei from the dorsolateral prefrontal cortex of individuals with MDD and healthy controls. Our analyses identified 26 distinct cellular clusters, and over 60% of these showed transcriptional differences between groups. Specifically, 96 genes were differentially expressed, the majority of which were downregulated. Convergent evidence from our analyses, including gene expression, differential correlation, and gene ontology implicated dysregulation of synaptic plasticity in the etiopathogenesis of MDD. Our results show that this high-resolution approach can reveal previously undetectable changes in specific cell types in the context of complex phenotypes and heterogeneous tissues.

## Introduction

Major depressive disorder (MDD) is a complex and heterogeneous disorder that affects an estimated 300 million people worldwide^1^. MDD can have serious implications, including death by suicide. The genetic factors underlying the risk for MDD have been investigated using different approaches, including genome-wide association studies^2^. Although some genetic associations have been detected, it has been difficult to identify specific and strong genetic correlates of the disease^3^.

The etiopathogenetic theory positing that MDD results from dysregulation of monoaminergic transmission, largely implicating the serotonergic and noradrenergic systems, has dominated the field for several decades. Recently, other factors have been associated with MDD, including glutamatergic and GABAergic transmission^4–7^, glial cell function, including astrocytic and oligodendrocytic contributions^7–14^, blood-brain barrier integrity^9^, and inflammation^15^. Thus, a wide variety of cell types found in the brain may contribute to the molecular changes underlying MDD.

In experiments conducted with cerebral tissue homogenates, the interpretation of differential gene expression is often complicated by the fact that the cellular composition of the sample is not uniform. Gene expression patterns in the brain are cell type specific, not only differentiating major classes of cells such as neuronal and glial cells, but even differentiating subtypes of glial cells and neurons^16,17^. Therefore, it is difficult to verify whether subtle molecular differences observed from tissue homogenates are explained by the disease state or by differences in cell type composition between samples^18,19^ and, just as gene expression patterns are cell type specific, it is likely that gene expression changes associated with MDD are also cell type specific. Recently developed techniques for high-throughput single-cell and single-nucleus RNA-sequencing provide a solution for addressing this inherent drawback to bulk tissue experiments^20–22^.

High-throughput single-cell RNA-sequencing (scRNA-seq) allows profiling of transcriptomes of individual cells by capturing the cells in nanolitre droplets using a microfluidic device and tagging every RNA molecule in a cell with a cell-specific barcode and a unique molecular identifier (UMI), all within the droplet. This method can also be extended to individual nuclei and yields comparable information^23,24^, allowing for analysis of frozen tissues, which are not amenable to the isolation of intact cells.

While there has been considerable interest in using single-cell datasets to gain insight into the processes underlying complex brain disorders^25,^, no direct comparison of single-cell human brain gene expression has yet been performed using high-throughput technologies.

We collected transcriptomic information on thousands of cells from 34 individuals, of whom half died during an episode of MDD, while the other half were psychiatrically healthy controls. We investigated differentially expressed genes across cell types and showed an overall enrichment for genes involved in synaptic plasticity, long-term synaptic potentiation, and synaptic organization. We also investigated the patterns of correlation of expression between differentially expressed genes across separate clusters.

Our results indicate that snRNA-seq can be successfully applied to complex psychiatric phenotypes to elucidate the role of specific cell types in pathology. The approach to snRNA-seq described here is an effective tool for interrogating subtle phenotypes with improved resolution in archived brain tissue from human brain banks, which are highly valuable but as yet, untapped resources for single cell transcriptomic research.

## Results

The human brain is comprised of many functionally unique cell types localized to specific brain regions and layers^22,23,26,27^, including some that have yet to be fully characterized. To assess the involvement of individual cell types in the pathophysiology of MDD, we examined nuclei from the dorsolateral prefrontal cortex (dlPFC), a region implicated in the pathology of major depressive disorder^6,8,11^. To assess a large number of nuclei, we used a droplet-based single nuclei method optimized for use with postmortem brain tissue. We assessed 78,886 nuclei from 34 brain samples, half from patients who died during an episode of MDD, and the other half from matched psychiatrically healthy individuals (Table 1, Supplementary Tables 1-3). The experimental design is depicted in Fig. 1a. On average, we sequenced to a depth of almost 200 million reads per sample (Supplementary Table 1). Glial cells have consistently been found to have fewer transcripts than neuronal cells^22,28^. We used custom filtering based on the distribution of UMIs detected (see Methods, Supplementary Fig. 1a-e, Supplementary Table 4) to recover a substantial number of glial cells. With an initial subset of 20 subjects, applying our custom filtering increased the total number of cells 1.8–fold but increased the number of non-neuronal cells by almost 6-fold (data not shown). More than 90% of the filtered cells had less than 5% mitochondrial reads, thus ensuring high quality data (Supplementary Fig. 1f). The average gene count across nuclei ranged from 2144 in neurons to 1144 genes in glia (Supplementary Table 5). UMI counts were approximately twice the gene count for all cell types, as expected for this level of sequencing depth (Supplementary Table 5). Between sample groups, there were no significant differences in the median gene count (t test p=0.12), median UMIs (t test p=0.14) and number of cells (t test=0.07) (Supplementary Table 1).

**Table 1:**
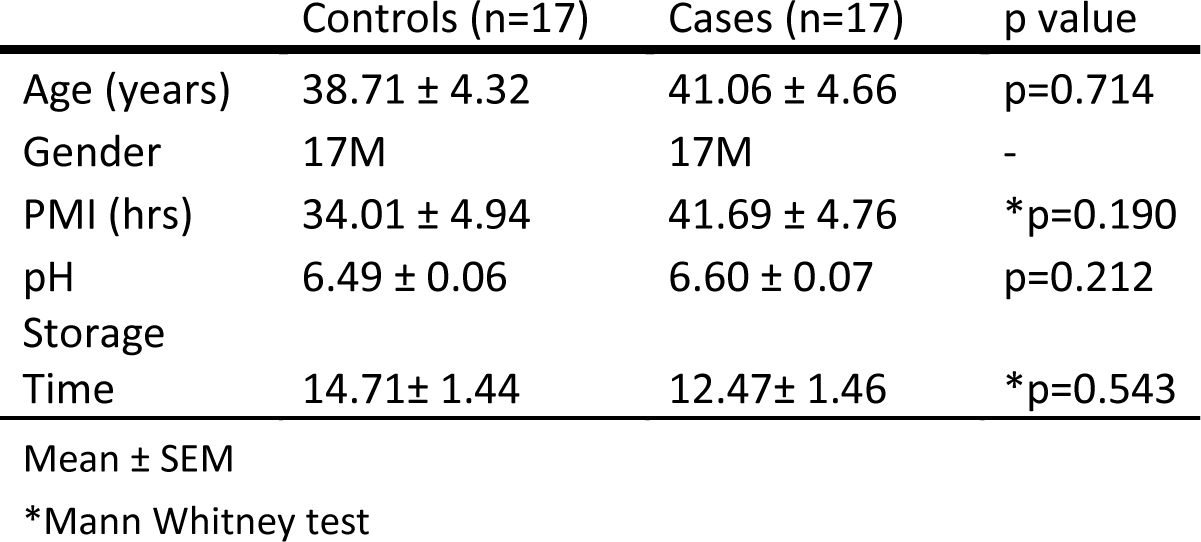
Sample information

**Figure 1:**
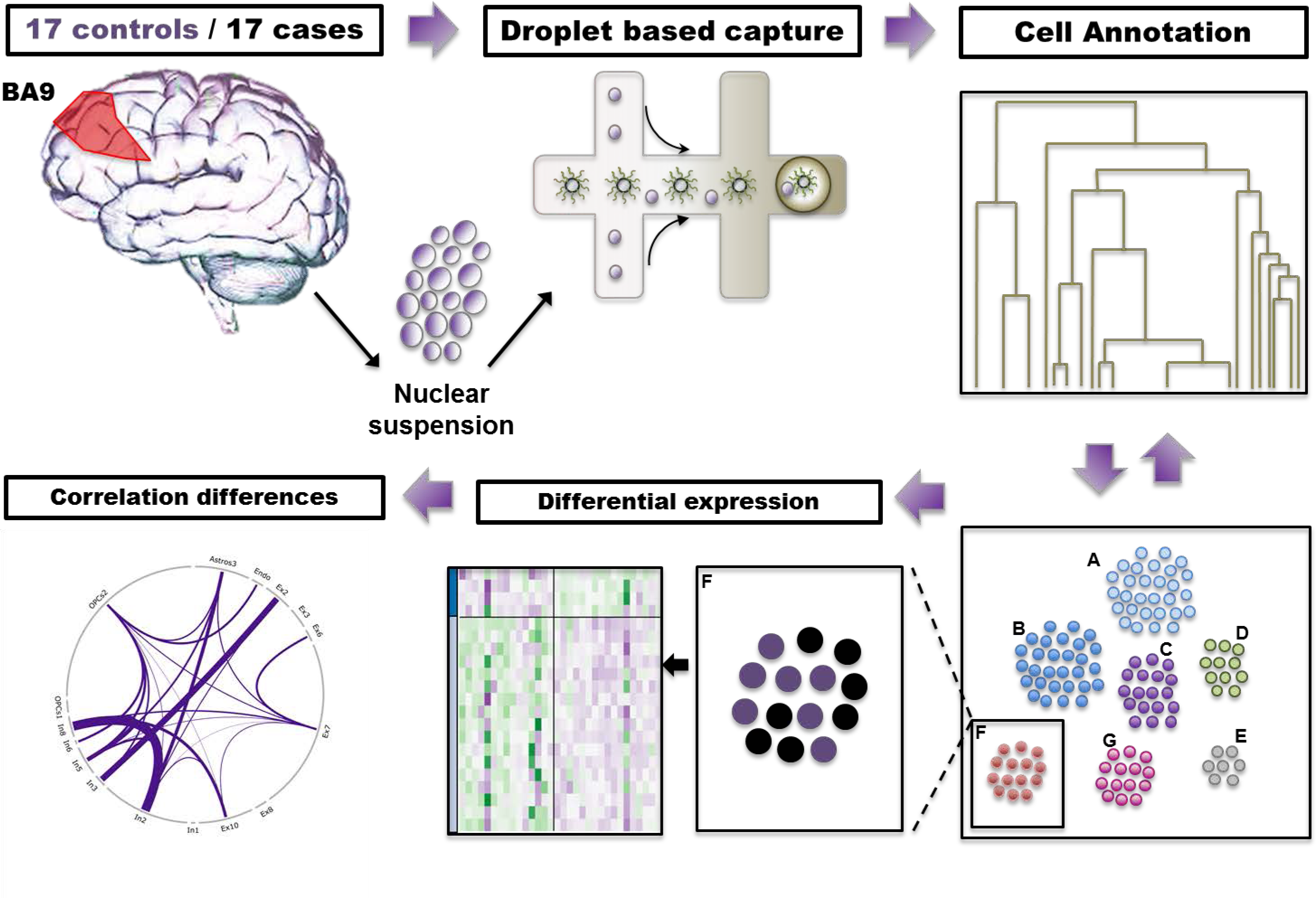
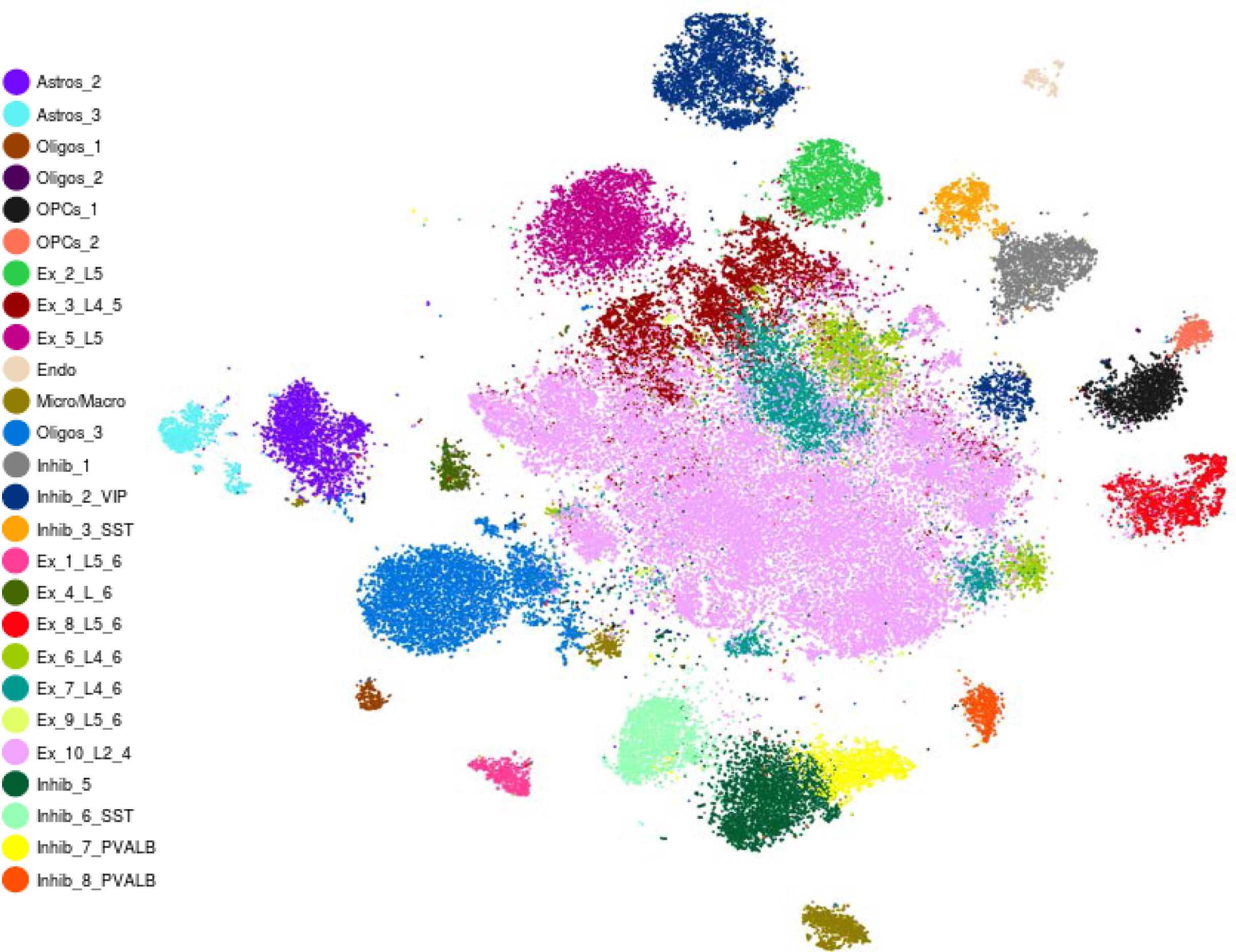
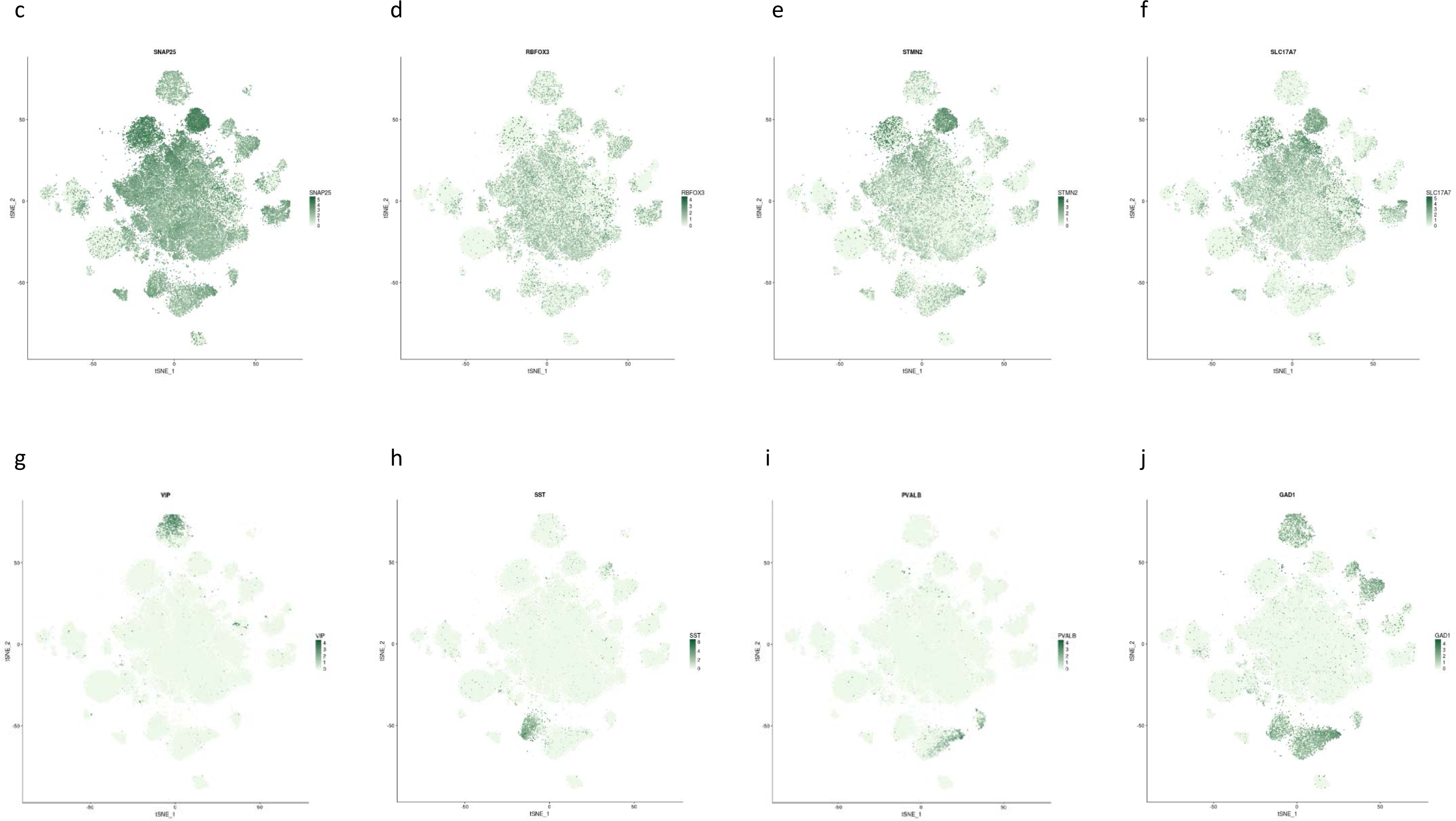
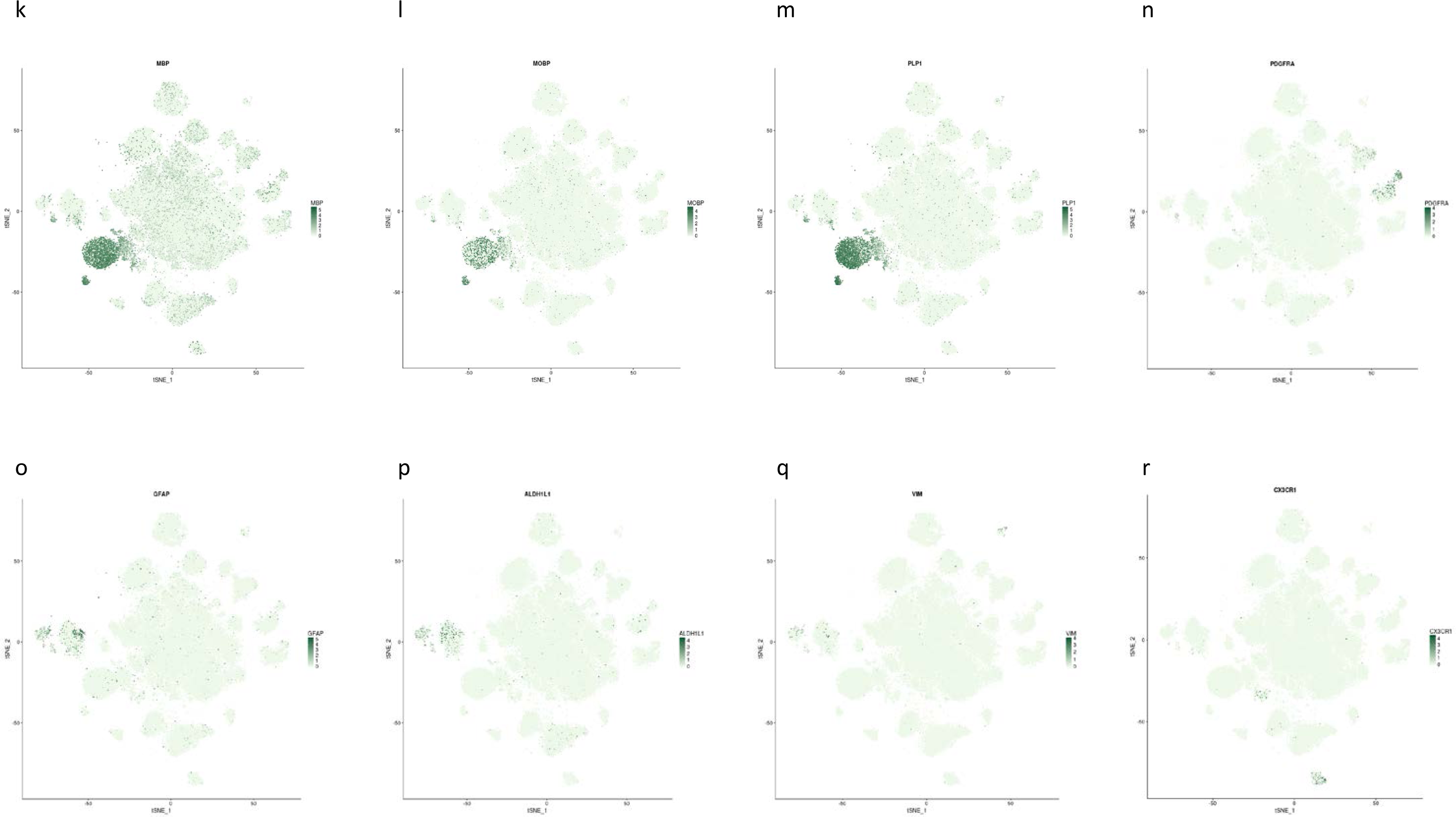
a) Schematic representation of experimental procedures. Nuclei were extracted from Brodmann area 9 (BA9) of the dlPFC of 17 cases and 17 controls, single nuclei were captured in droplets for RNA-seq. Unsupervised clustering and cell type annotation were followed by differential expression analysis between cases and controls within each cluster. b) TSNE plot depicting the ~73,000 cells in 26 clusters identified after strict quality control of initial clusters. Includes 2 astrocytic, 3 oligodendrocytic, 2 oligodendrocyte precursor, 1 microglial, 1 endothelial, 10 excitatory neuronal and 7 inhibitory neuronal clusters. The majority of cells are present in excitatory clusters. Individual TSNE plots representing the expression of various neuronal (c-j) and non-neuronal (k-r) cell type marker in a given cluster.

### Unsupervised clustering identified 26 unique cell types in the dlPFC

In order to identify different cell types present in the brain samples, we applied unsupervised graph-based clustering^29^ using the first 50 principal components derived from the 2135 most variable genes across individual cells (see Methods, Supplementary Fig. 2a-b). This initially resulted in 30 distinct cell types with the majority (47,461) of cells belonging to excitatory clusters, as expected^30^, based on an initial annotation (Supplementary Fig. 2c-d). We then reprocessed all excitatory clusters whose average gene expression profiles were mutually highly correlated (R>0.95), this included 7 clusters of ~40,000 cells. Two smaller excitatory clusters not sufficiently correlated with these 7 were not re-clustered. These ~40,000 cells were reprocessed using similar parameters for clustering as the whole dataset (with slight differences, see Methods) producing a refined sub-clustering of excitatory cell types (Supplementary Fig. 3a-c, Supplementary Table 6). Finally, the clusters were manually curated to eliminate potential biases; for example, clusters were removed if mainly one sample contributed to the cells contained within the cluster (Supplementary Tables 7-10, Supplementary Fig. 4a-c). After stringent quality control (see Methods), we identified 26 unique clusters (Fig. 1b). Each cluster was annotated using a combination of known cell markers including broad cell markers to identify neurons in general (*SNAP25, RBFO3, STMN2*), excitatory cells (*SATB2, SCL17A7, SLC17A6*), inhibitory cells (*GAD1, GAD2, SLC32A1*) (Fig. 1c-j), and non-neuronal cells, including astrocytes (*GLUL, SOX9, AQP4, GJA1, NDRG2*), oligodendrocytes (*PLP1, MAG, MOG, MOBP, MBP*), oligodendrocyte precursor cells (OPCs) (*PDGFRA, PCDH15, OLIG1, OLIG2*), endothelial cells (*CLDN5, VTN*), and macrophages/microglia (*SPI, MRC1, TMEM119, CX3CR1*) (Fig. 1k-r). Gene expression patterns specific to cell type clusters were visualised using DotPlots (Fig. 2a), average expression and median expression heatmaps (Supplementary Fig. 5a-b) and violin plots (Fig. 2b-e) to form a consensus for annotation.

**Figure 2:**
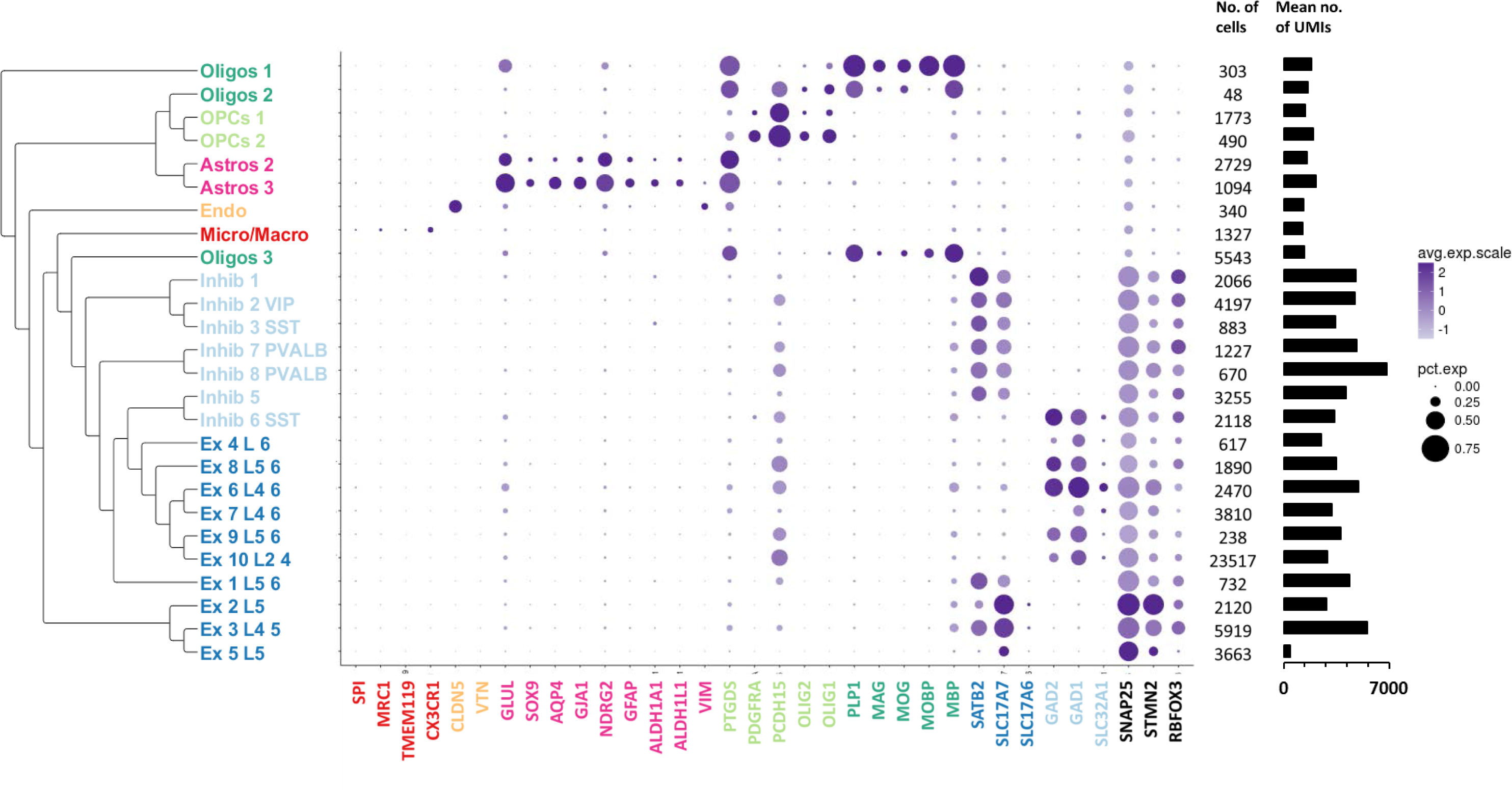
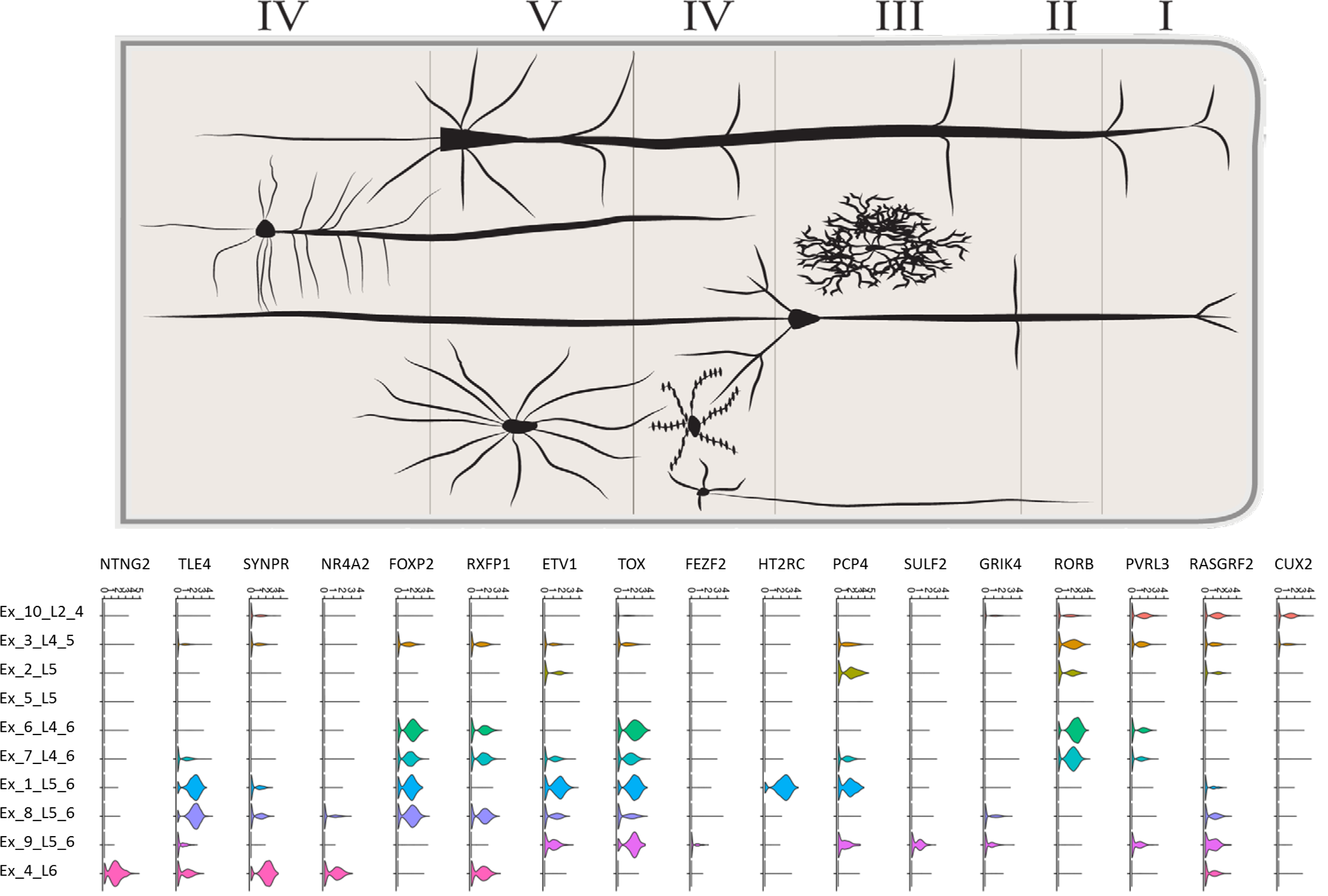
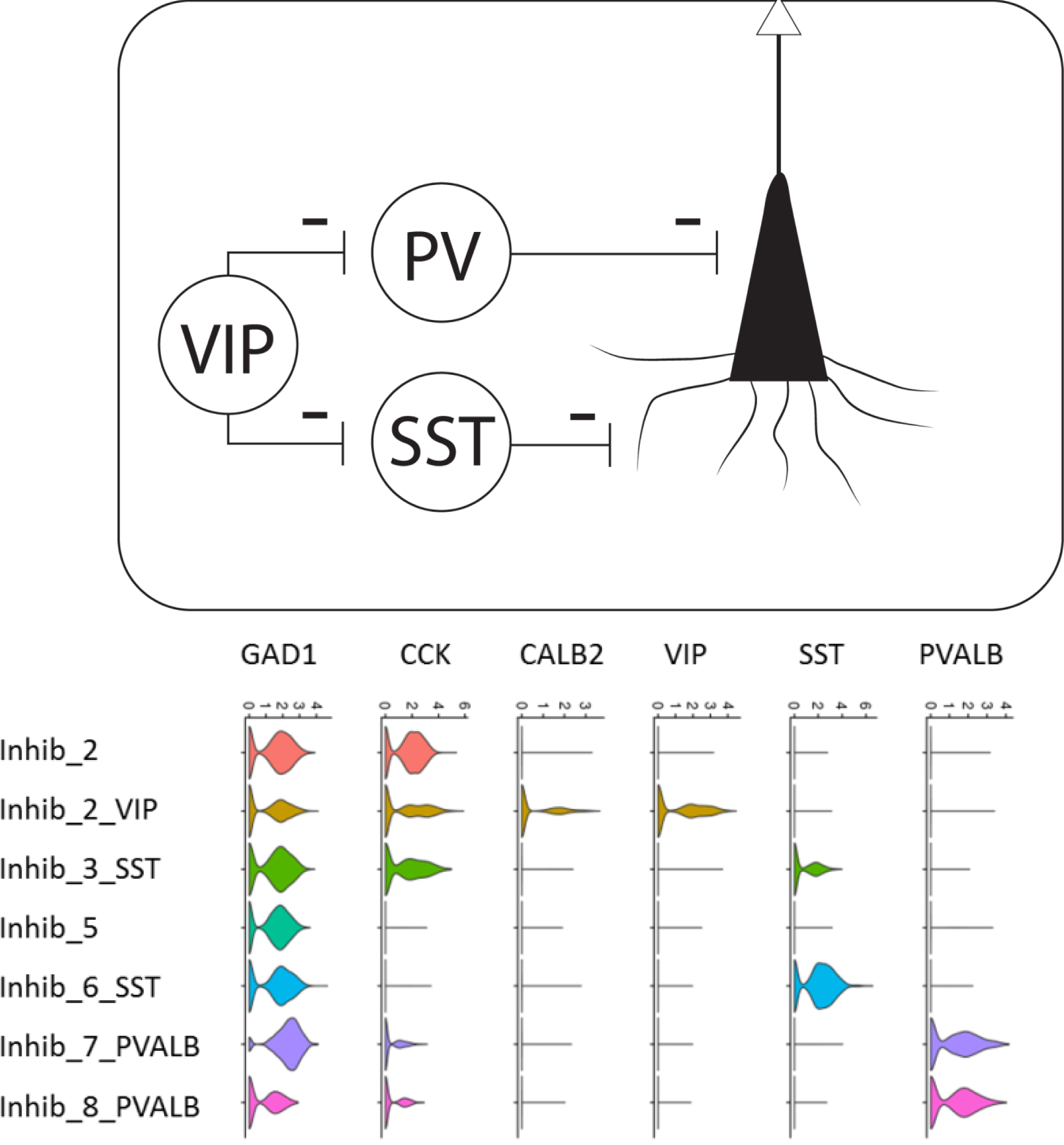
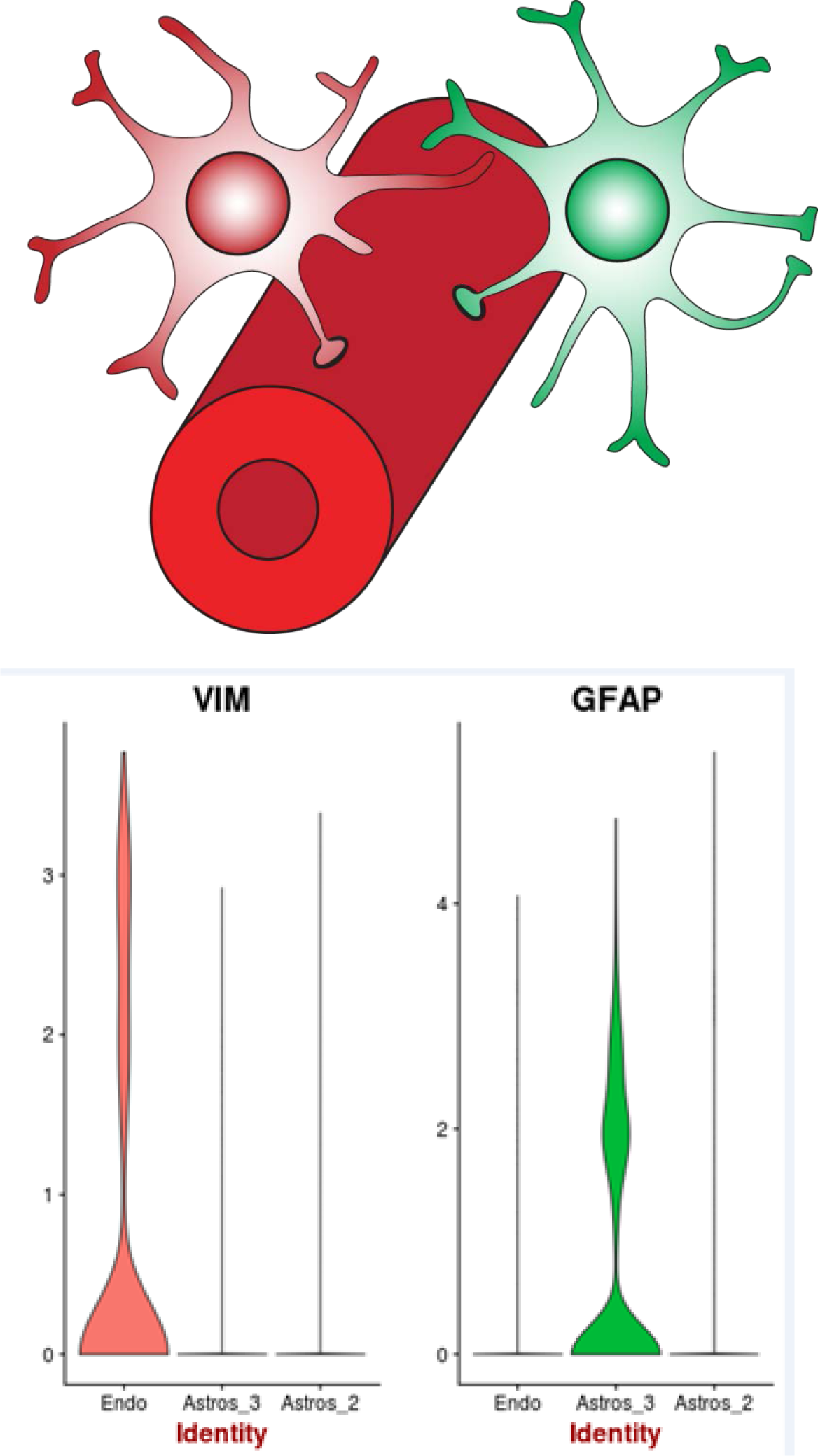
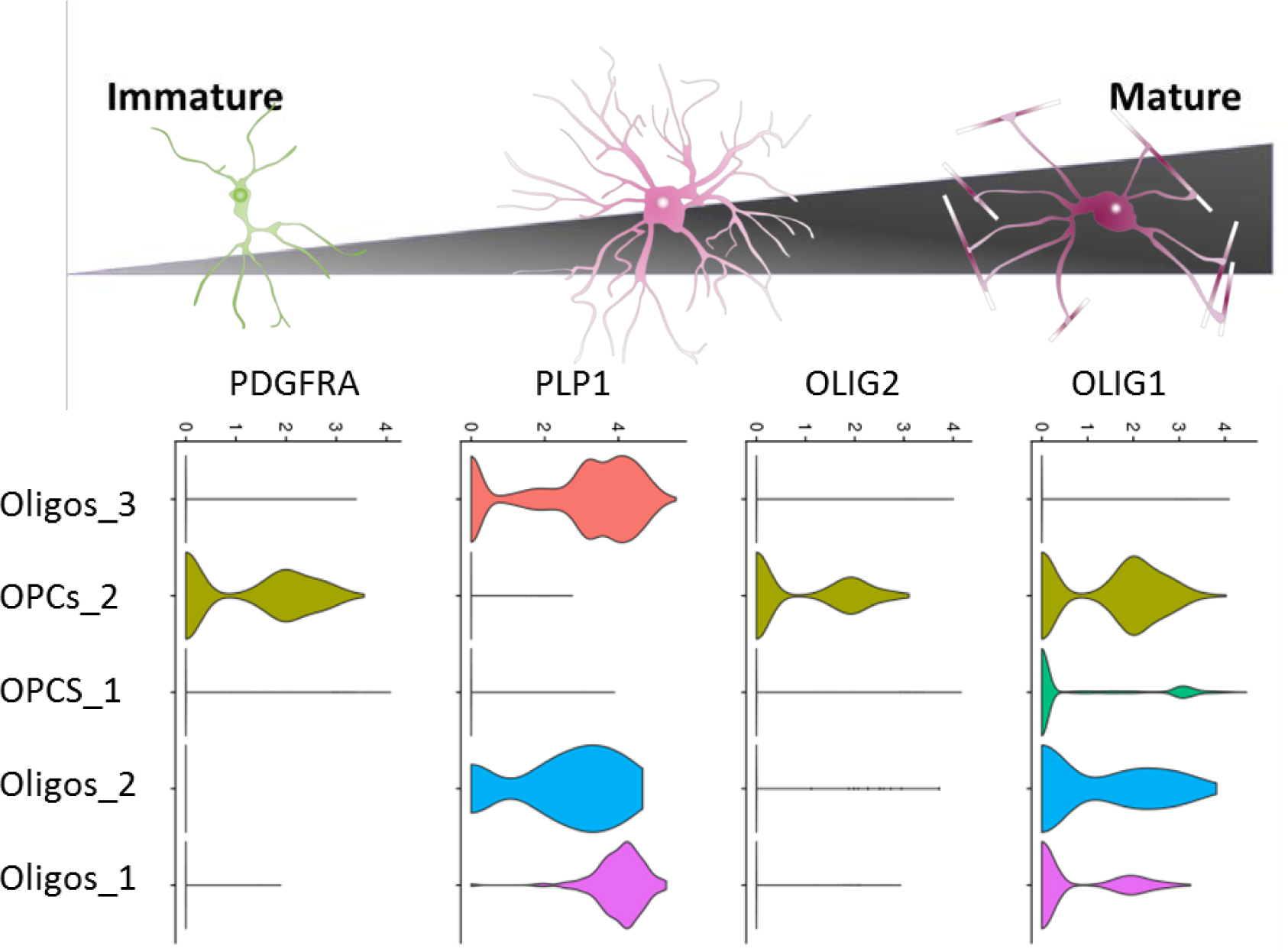
a) Cell type annotation was performed based on expression of well-established marker genes. (Left) Dendrogram representing relationship between identified cell clusters based on gene expression. (Middle) DotPlot depicting expression of known marker genes in the 26 clusters of interest. Marker genes are colour coded according to the cell type in which they should be detected (e.g.: red for SPI, which is expected to be microglial). The size of the dots represents the proportion of cells expressing the gene whereas the colour intensity represents the average expression level. (Right) The list of numbers gives the size of each cluster and the bar plot depicts the mean number of UMIs per cell in each cluster. Overall, non-neuronal cell types show lower mean number of UMIs. b) Cortical layer specific markers varied in expression within the excitatory neuronal clusters produced after sub-clustering of initial clusters. The schematic of cortical layers can be used to orient the marker genes to the appropriate layer. The violin plots depict the expression per cluster of layer specific marker genes going from the more superficial layers on the right (starting from *CUX2*) to the deeper layers on the left (ending at *NTNG2*). Excitatory clusters were annotated with their approximate layer-specific identities based on the expression pattern observed. c) Refined inhibitory cluster identification. Known classes of inhibitory neurons are identifiable based on the expression pattern of peptide genes (*VIP, SST, CCK*) and calcium binding protein genes (*PVALB*). d) Astrocyte and non-neuronal cells. Higher GFAP expression in Astros_3 than Astros_2 may reflect their reactive state. We were unable to detect vimentin (*VIM*) positive astrocytes, although *VIM* expression was detected in endothelial cells as expected. e) Cells belonging to the oligodendrocyte lineage.

**Figure 3.**
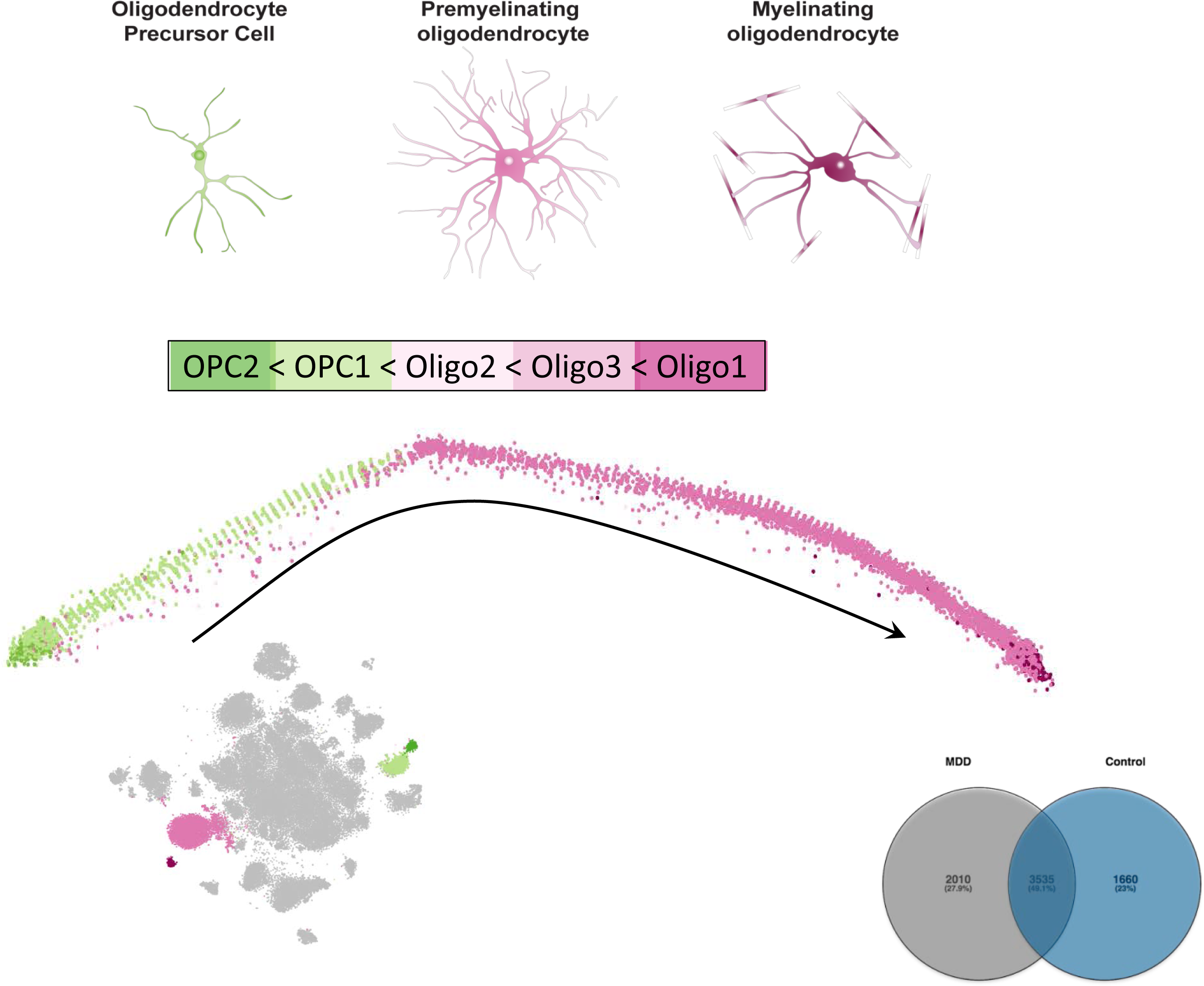
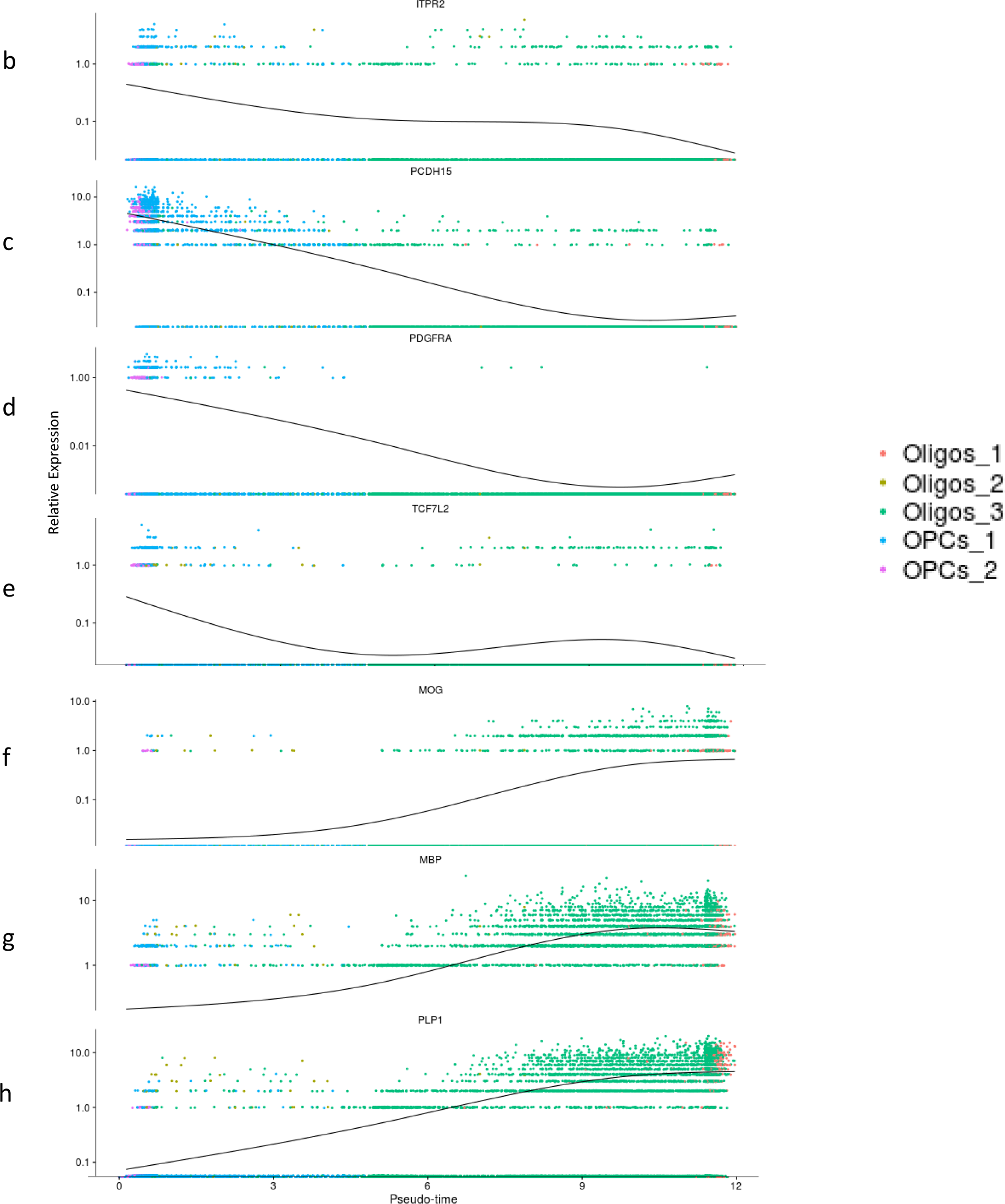
Pseudotime trajectory. a) (Left) Oligodendrocyte lineage cells from 5 clusters were analysed to produce a pseudotime trajectory to gauge their developmental stages. Going from left to right along the trajectory we see a progression of immature and mature oligodendrocyte precursor cells followed by immature and mature oligodendrocytes in the order shown. Inset provides the positions of the clusters in the original TSNE plot. (Right) The diagram shows the number of genes that changed in expression with pseudotime separately in cases and controls. While 3535 gens were common to both groups, 2010 were only associated with pseudotime in cases, and 1660 were only associated with pseudotime in controls. Expression across pseudotime of (b-d) genes known to be highly expressed in OPCs or immature oligodendrocytes (e) transitionary, or (f-h) highly expressed in mature oligodendrocytes.

**Figure 4:**
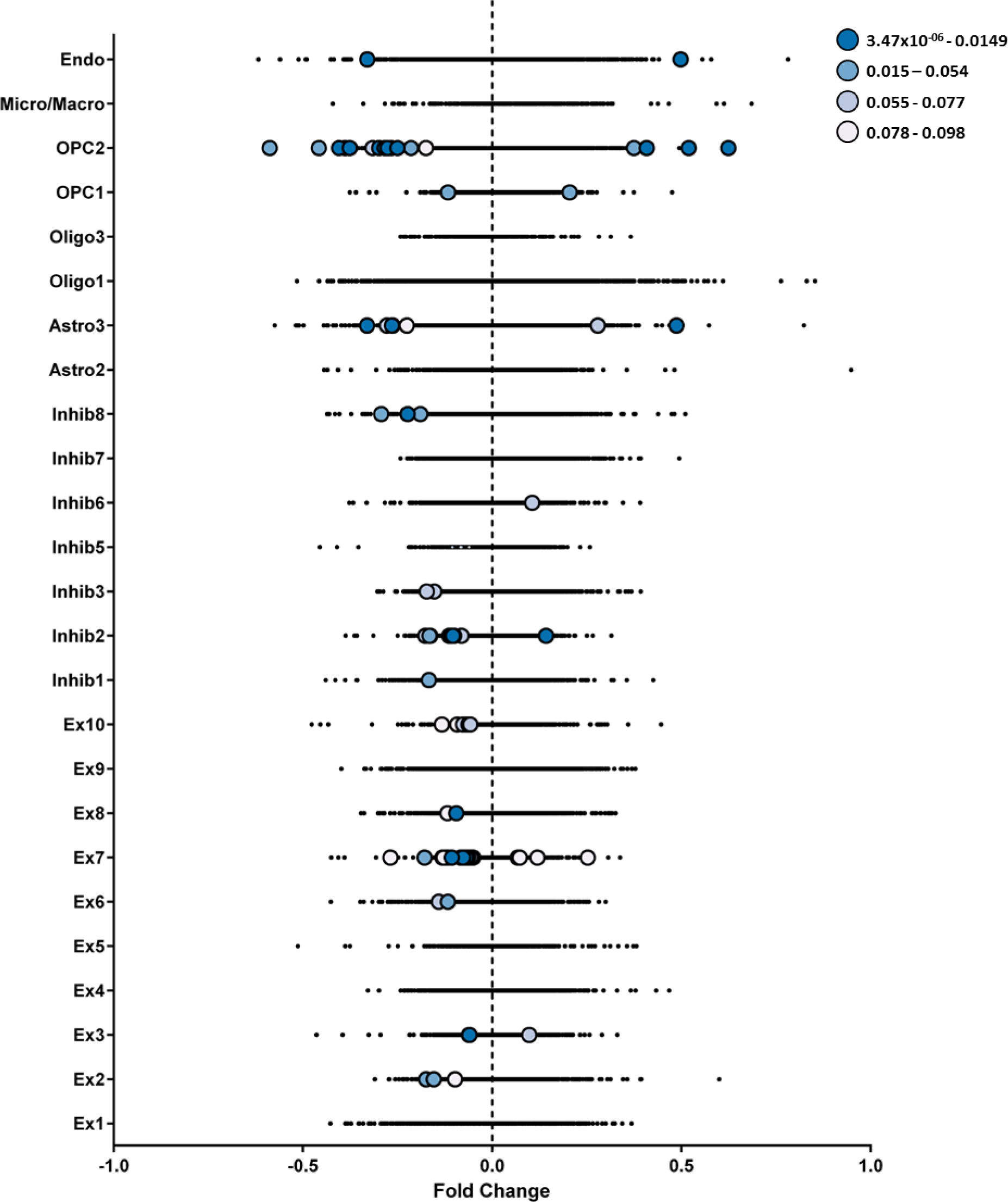
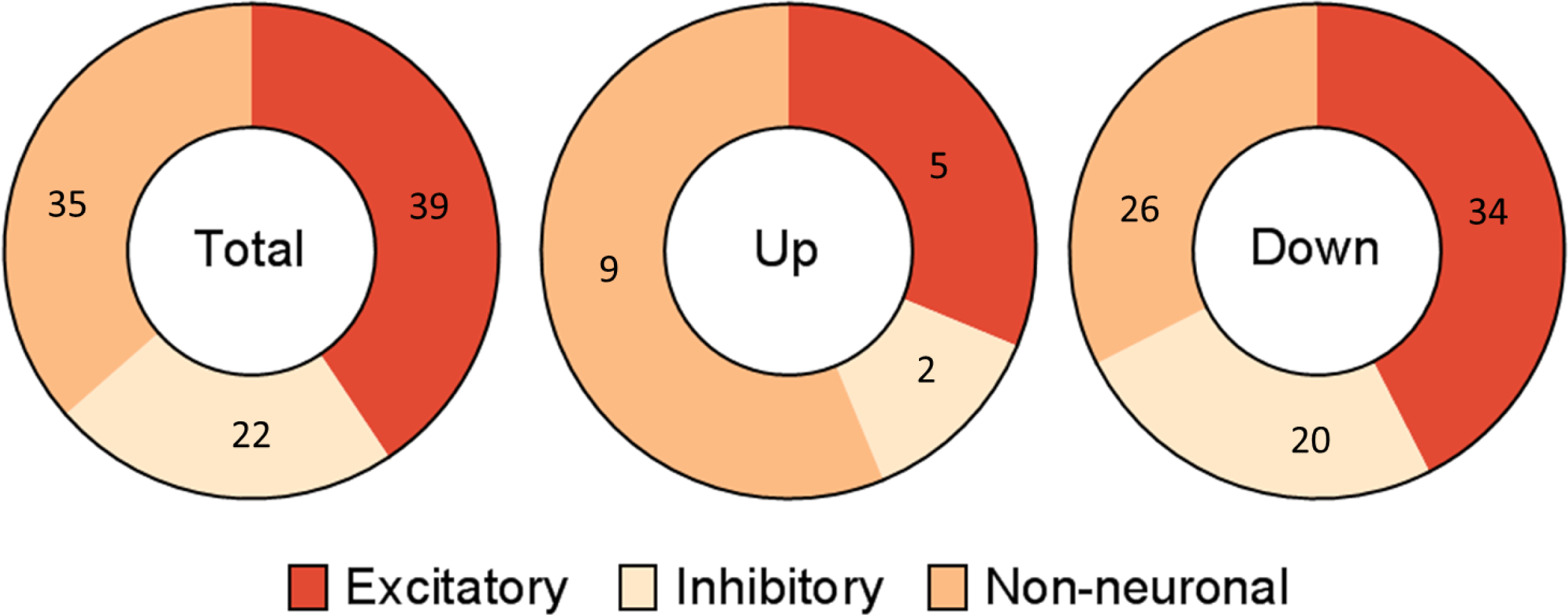
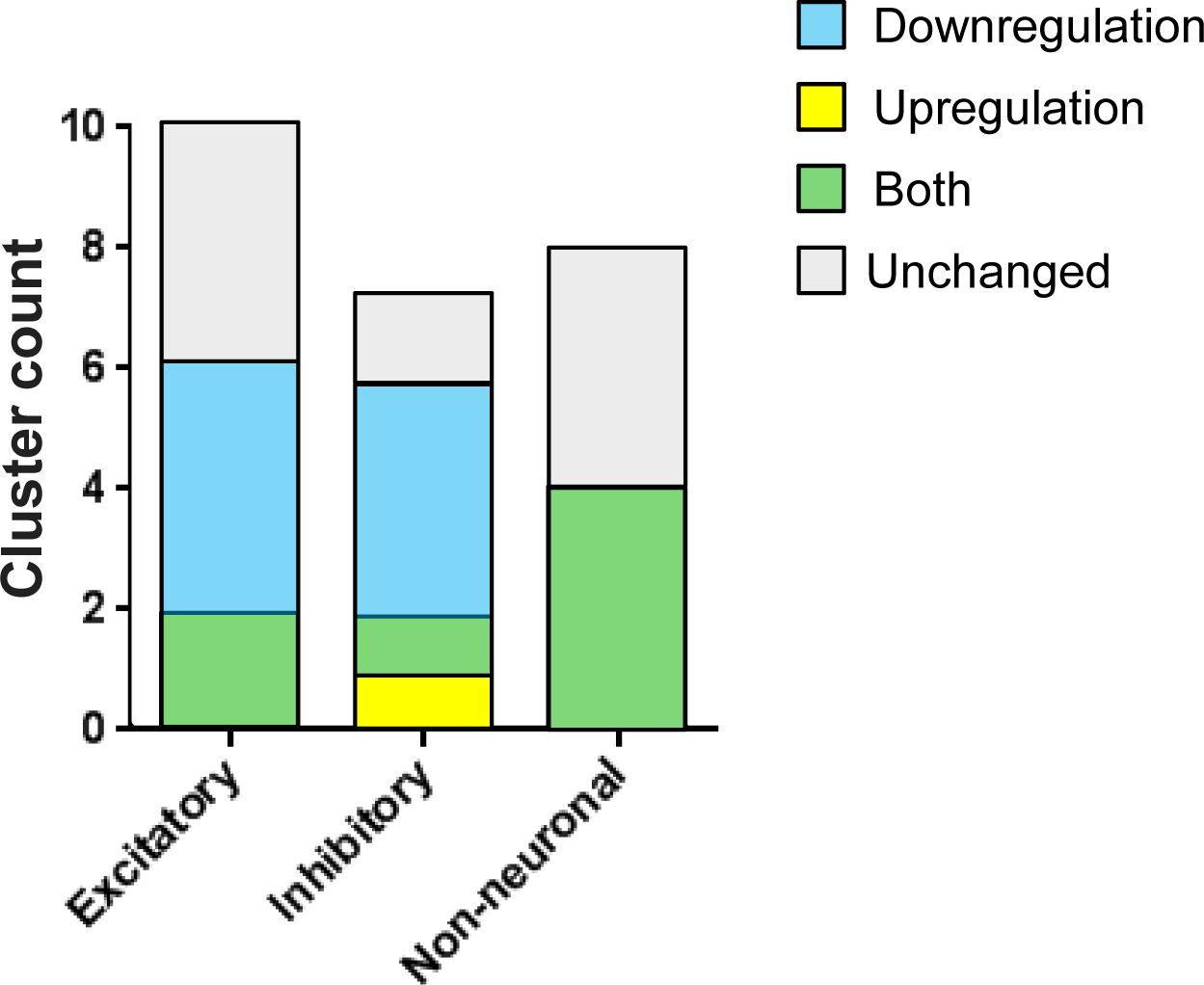
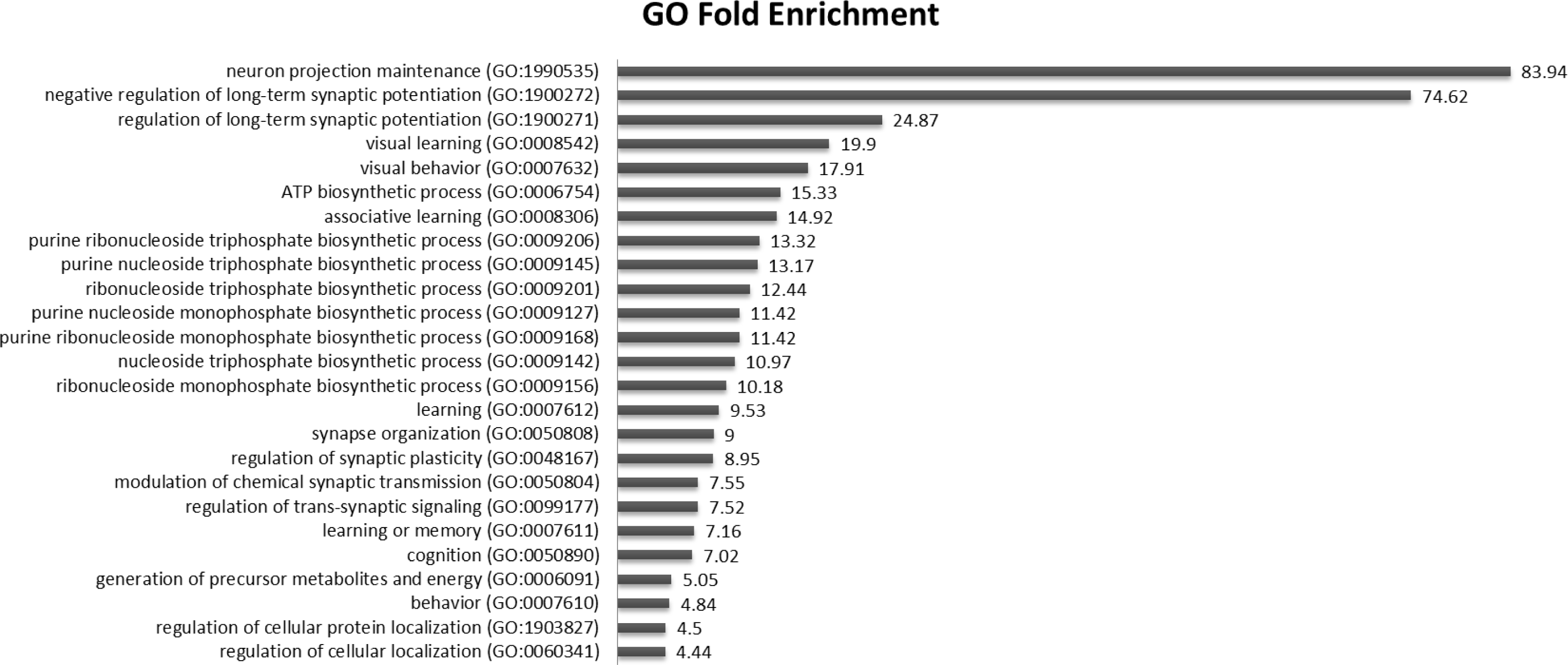
a) For each cluster the percentage change in expression between cases and controls of all detected genes are plotted with decreased expression to the left of the midline and increased expression of the right. Ninety-six significantly changed genes (of which 16 were upregulated and 80 were downegulated) are marked in colour, based on their corrected FDRs as shown in the legend (light to dark blue corresponds to higher to lower FDRs). Sixteen out of the 26 clusters contained significantly differentially expressed genes b) Contribution of different cell type clusters to differentially expressed genes is depicted in pie charts. While the proportion of total differentially expressed and downregulated genes contributed by the different cell type clusters were relatively similar, non-neuronal clusters contributed a higher proportion of genes upregulated in MDD cases. c) Number of clusters in each broad category showing up and downregulated genes in MDD cases. While some excitatory and inhibitory clusters showed only downregulation, all dysregulated non-neuronal clusters contained both up and downregulate genes. A larger proportion (7/8) inhibitory clusters showed dysregulation compared to excitatory (6/10) or non-neuronal (4/8) clusters. There was just one cluster (inhibitory), which showed only upregulation. d) Top 25 gene ontology terms associated with the 96 differentially regulated genes.

**Figure 5:**
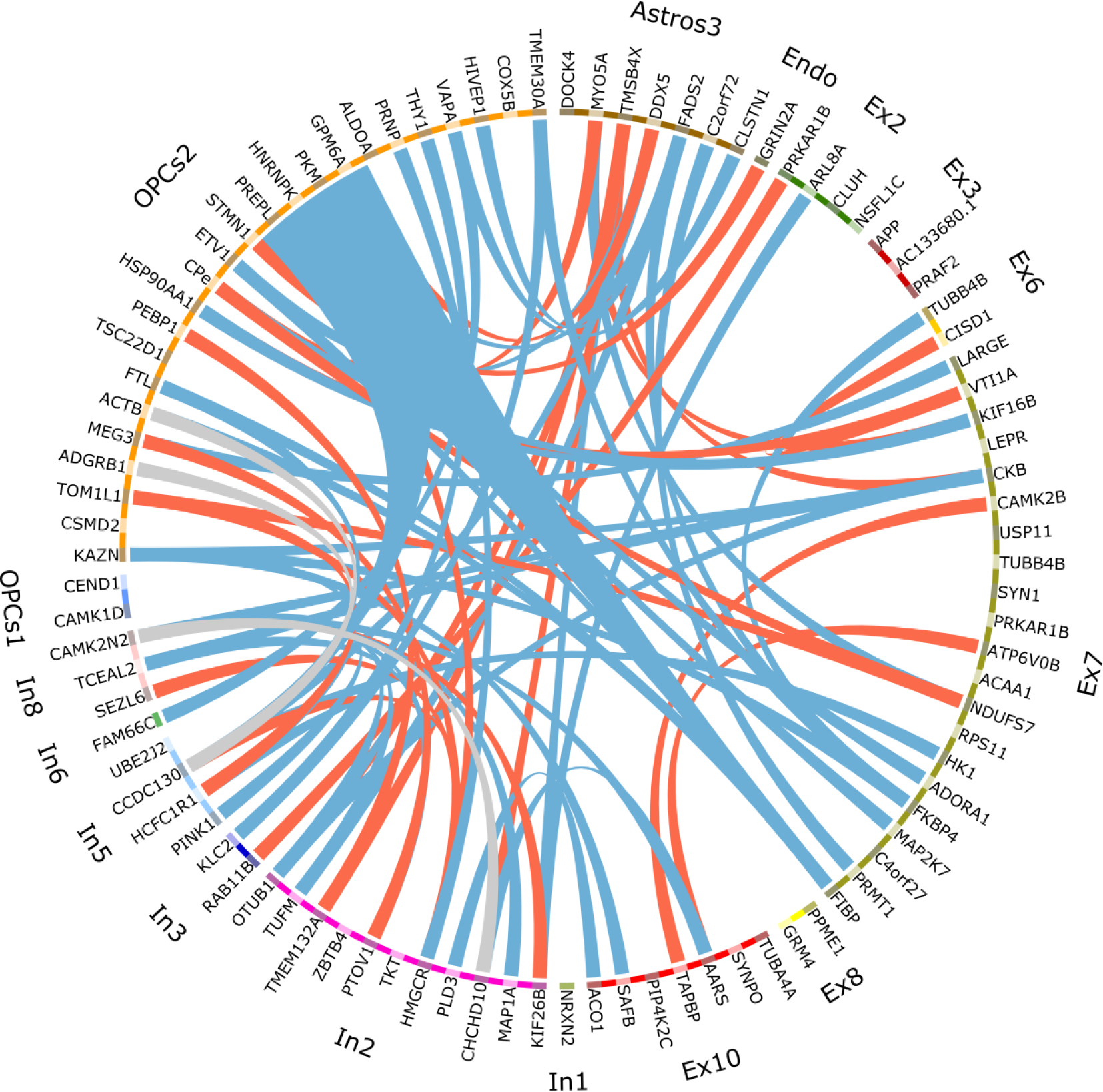
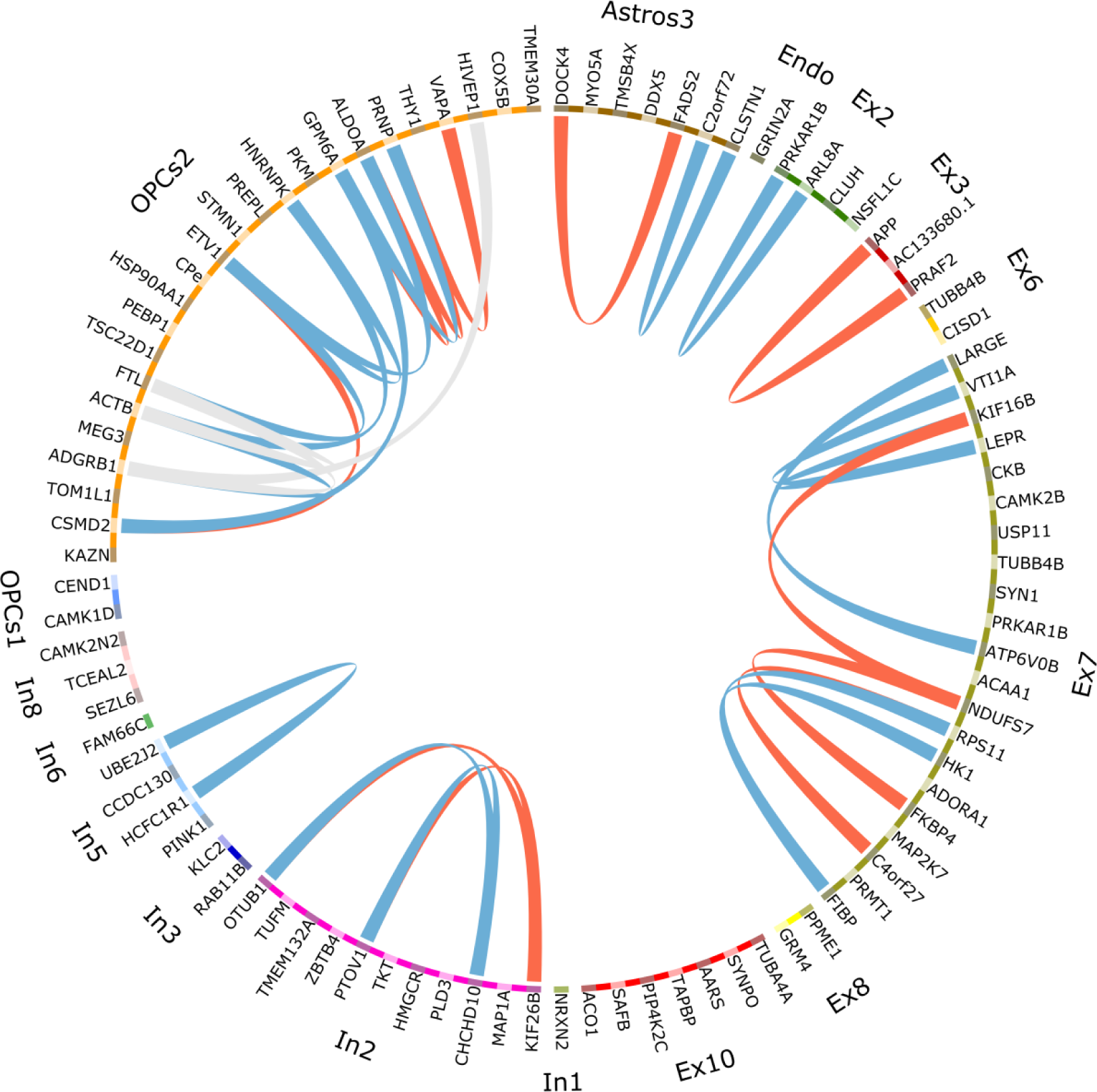
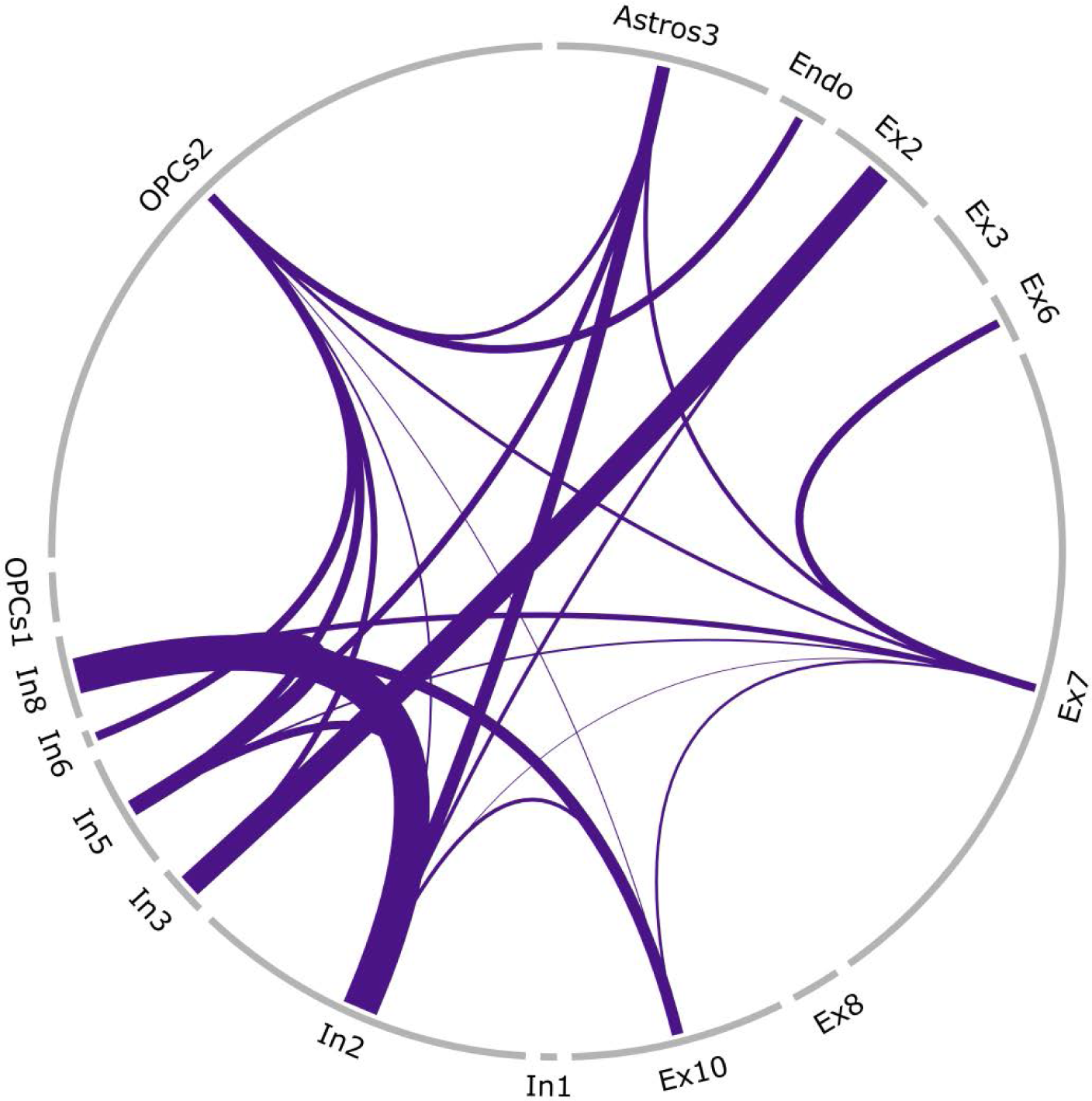
Significant z-scores for pairs of 95 (one dropped) differentially expressed genes in the 16 dysregulated clusters are represented in a Circos plot. a) Only the z-score for pairs of genes coming from two different clusters are shown here (between cluster z-scores). The gene names and the clusters that contain them are labelled outside the circle. Lines connecting genes represent that the correlation of gene expression for that pair of genes was significantly different between MDD cases and controls. Blue lines indicate positive correlation in cases and negative in controls, red lines indicate positive for control and negative for cases, and grey lines indicate the same direction of correlation but different strengths. b) As in (a) but for pairs of differentially expressed genes in the same cluster. c) Circos plot depicting weighted (see Methods) lines showing the overall level of correlational differences between different clusters. In brief, the thicker lines connect clusters whose correlations change more strongly between MDD cases and controls.

### Refining cell types

The clusters generated from our data are consistent with previously reported gene expression patterns that vary within cell types (Supplementary Fig. 6), though our considerably larger sample set allowed us to produce more unique clusters than previously observed^22^.

Gene expression patterns previously linked to specific cortical layers (see Methods) coincide with our clustering of excitatory cells. In Fig. 2b, the genes are arranged from left to right in order of their expression across the cortical layers (from the layer VI to layer II). There is a gradient of expression of these genes across the excitatory clusters. For example, clusters Ex1, Ex4, and Ex7-9 had high expression of *TLE4* (layer VI specific). Ex1, Ex8, and Ex9 showed concurrent expression of layer V/VI markers such as *TOX*. Ex6 and Ex7 additionally showed expression of the layer IV specific gene *RORB. HTR2C*, which is specific to a subset of layer V neurons, was prominent in Ex1 alone. *PCP4*, which is also layer V specific, was present in Ex1-3, Ex7, and Ex9. Superficial layer (I-III) markers such as *CUX2* and *RASGRF2* were mainly seen in the large cluster Ex10. Likewise, inhibitory cell types demonstrated subtype specific gene expression patterns. For example, In7 was classified as inhibitory parvalbumin because it expressed *GAD1* and *PVALB*, and lacked *VIP* and *SST* (Fig. 2c). Multiple astrocytic clusters were also identified, and while the typical sub-classification of astrocytes is based on their morphology within grey or white matter^31^, we used only grey matter for these samples. As such, based on the higher percentage of *GFAP* expression in Astros_3 (38%) compared to Astros_2 (21%), we expect Astros_3 to represent reactive astrocytes^32^ (Fig. 2d, Supplementary Table 11).

### Oligodendrocyte cell lineage

We identified five unique cell clusters that fell into the oligodendrocyte lineage (OL), including two that we classified as OPCs (Fig 2e). OPCs express a characteristic set of markers such as PDGFRA^33^ and PCDH15^34^, which decline as these cells mature into oligodendrocytes, whereas other lineage markers like, OLIG2 or SOX10, are present in both mature and immature cells^33^. Given these developmental stage specific markers it is possible to plot a pseudotime trajectory^35^ using gene expression for OPC1, OPC2, Oligos1, Oligos2 and Oligos3. The result indicates that OPC2 are the youngest cells within the dataset followed by OPC1, then Oligos2 and Oligos3, with Oligos1 being the most mature (Fig. 3a). The expression of thousands of genes varied according to pseudotime (q<0.01), but approximately half of the associations were observed in both cases and controls (Fig. 3a, right). Among the genes that are uniquely associated with pseudotime in cases, there was a 2.7–fold enrichment of apoptosis signalling (FDR p<9.01×10^-3^)^36^, while no functional enrichment was observed in controls. To assess the individual profiles of important developmental gene markers, we plotted their expression across pseudotime (Fig. 3b-h), revealing their expected pattern of expression.

### Within cluster gene expression differences associated with depression

We set out to assess gene expression differences between cases and controls within each cluster. However, one limitation of droplet based single-cell technology is the possibility of capturing doublet or multiplet nuclei in a given reaction. These represent a potential confounding factor when assessing differential gene expression between groups. We therefore attempted to eliminate them from the dataset by calculating the correlation of each cell to the median expression value of its assigned cluster (See Methods, Supplementary Fig. 7). Cells with low correlation were removed. We also excluded any genes expressed in less than 10% of the cells in that cluster. Using only these purified clusters and filtered genes, we performed a differential gene expression analysis (Supplementary Tables 12-36). Olig2 was excluded from differential expression analysis because it contained only 48 cells.

A total of 96 genes (FDR <0.1) were differentially expressed in 16 of the 25 clusters analyzed (Fig. 4a) and 45 of those remained significant at FDR<0.05 (12 of 25 clusters). The majority, 80 genes (83%), were downregulated, in line with findings from previous transcriptomic studies in MDD^3,5,7^. While the differential expression analysis treated each cell as a sample, per subject contributions to the differential analysis were visualized using heatmaps of average gene expression (Supplementary Fig. 8a-p) to assess biases in sample contributions. Thirty-nine of the 96 differentially expressed genes were found in excitatory cell clusters and, of those, 34 were downregulated (Fig. 4b). Certain neuronal clusters contained both upregulated and downregulated genes, however it was more common for clusters to have only downregulated genes. Non-neuronal clusters tended to have both up- and downregulated genes (Fig. 4c). There were two instances of the same gene being differentially expressed in separate clusters: *PRKAR1B* showed decreased expression in excitatory clusters Ex7 (FDR=0.087, FC=0.87) and Ex2 (FDR=0.047, FC=0.82) and *TUBB4B* in excitatory clusters Ex7 (FDR=0.079, FC=0.87) and Ex6 (FDR=0.073, FC=0.86).

There were strong enrichments of Gene Ontology terms for *neuron projection maintenance* (84-fold enrichment; FDR=0.011) and *negative regulation of long-term synaptic potentiation* (75-fold enrichment; FDR=0.012). Both of these terms are hierarchically related with the more general term *regulation of synaptic plasticity*, also enriched in the set of 96 genes (9-fold enrichment, FDR=0.012) (Fig. 4d).

### Between and within cluster correlations as indications of how cells interact in MDD

To assess how interactions between cells might contribute to psychopathology, we assessed the correlation of differentially expressed genes between clusters. Average expression per subject for each of the differentially expressed genes was calculated in each of the 16 clusters. A correlation coefficient for each pair of genes was independently calculated for cases and controls and transformed into a Fisher z-score for comparison between groups (see Methods). Any z-score with p<0.01 was retained. The significant correlation differences between clusters are represented in Fig. 5a. Any differential correlations resulting from a magnitude difference in correlation coefficients are represented in grey as they are believed to demonstrate consistency in the biological function between groups. A positive z-value (blue) arises from gene pair correlations that are positive in cases and negative in controls, whereas a negative z-value (orange) arises from the opposite combination. Of equal interest, we examined clusters that showed high levels of within-cluster correlation differences. As expected, OPC2 and Ex7 showed the highest number of changes in gene relationships within clusters given they have the highest number of differentially regulated genes. Interestingly, the Ex3 cluster has no between cluster relationship but shows a within cluster change (Fig. 5b). We used Fisher’s method to assess the overall correlation differences between clusters (Fig. 5c). This shows the strongest difference in the correlations between In2 and In8.

## Discussion

The complex arrangement of numerous cell types in cortical cytoarchitectonics makes it difficult to tease out the respective implication of these cells in MDD and other brain illnesses. It is only with the advent of single-cell technology that we are beginning to understand the total number of cellular subpopulations that exist in the brain^17^. Given the complexity of psychiatric disorders such as MDD, and the absence of consistent, salient genetic contributions, disentangling the role of each cell type in the brain is of great importance and will require the level of resolution we have achieved here.

Compared to previous single-nucleus PFC transcriptomic studies, our larger data set allowed us to resolve a greater diversity of excitatory clusters than both droplet-based^22^ (10 vs 2) and full length snRNA-seq protocols^26^ (10 vs 8). Custom filtering increased non-neuronal cell content, allowing greater resolution of glial subtypes, including multiple astrocytic, oligodendrocytic, and OPC clusters. For example, this resolution enabled us to pinpoint changes specific to OPCs but not oligodendrocytes, and changes selective to one subset of astrocytic cells. The same principle extends to neuronal cell types.

Differential gene expression analysis suggests synaptic function alterations in depression, including *synaptic plasticity*, *regulation of long-term synaptic potentiation*, *synaptic organization* and more broadly, learning, memory and cognition. Genes such as *APP* (Ex3), *PRKAR1B* (Ex2/7), and *PRNP* (OPC2), were consistently associated with a number of the most enriched terms related to synaptic function and cognition.

Interestingly, *APP* was found to be differentially correlated with *PRAF2*, both of which were found to be dysregulated within Ex3 (IV/V). There was a strong negative correlation (R=-0.93) in MDD cases but a moderate association in controls (R=0.25). The products of these genes interact directly^37^ and both are important for synaptic function^38,39^. This is suggestive of synaptic dysregulation in layer V pyramidal cells in the PFC of MDD cases. Given that layer V is primarily composed of projection neurons^40^, this may implicate other brain regions involved in mood and emotions, including the limbic system.

We found that *PRKAR1B* (encoding protein kinase cAMP-dependent type I regulatory subunit beta) was decreased in MDD cases in 2 separate clusters Ex7 (IV-VI) by 12% and in Ex2 (V) by 15%. PRKAR1B is involved in the cAMP second messenger-signalling pathway and in dopamine receptor signaling^41,42^. Dopamine (DA) is an important modulator of synaptic plasticity in the PFC^43,44^. Dopaminergic afferents from the ventral tegmental area (VTA), project primarily to layer V in the PFC^17^, and although DA is often overlooked in MDD, recent evidence points to altered dopaminergic signal transduction in depression ^45^.

Additionally, *PRKAR1B,* from Ex2, was differentially correlated with *RAB11B* from In3 (SST). *RAB11B* (encoding for a Ras-related protein) is critical in vesicle transport and recycling^46^, in particular, trafficking and recycling of monoamine transporters^47^. It is demonstrably involved in dopamine transporter (DAT) recycling^48^. Both *PRKAR1B* and *RAB11B* were decreased in MDD cases and there was a strong negative correlation between these two genes (R=-0.83) in MDD cases, but a strong positive correlation in controls (R=0.59). It is possible that in healthy individuals these proteins work synergistically to regulate dopaminergic signal transduction. In this study, we found a reduction of *RAB11B* and *PRKAR1B* which may disrupt DAT trafficking and inhibit DA mediated second messenger signalling, respectively. Together, these changes may indicate impaired DA signal transduction and DA-related synaptic plasticity, although further empirical evidence is required to support this hypothesis.

In addition to being associated with mediating synaptic plasticity^49,50^, the <u>pr</u>io<u>n</u> <u>p</u>rotein gene (*PRNP*) was strongly decreased (28%; FDR=0.038) in the OPC2 cluster, the least developed cells identified in this study. The absence of *PRNP* has been associated with an increased number of undifferentiated oligodendrocytes, and OPCs that do not mature into oligodendrocytes are thought to be eliminated by apoptosis^51^.

Indeed, we uncovered evidence suggesting that MDD may fundamentally modify the developmental trajectory of oligodendrocytes. The gene sets associated with developmental trajectory differed significantly between MDD cases and controls. We also observed an enrichment of apoptosis-related genes in cases but not in controls. This is in line with both the described effects of decreased *PRNP* in OPCs^51^, and evidence of a loss of mature adult oligodendrocytes in animal models of depression and anxiety^52^. Interestingly, there is evidence suggesting that half of the OPCs (NG2^+^) in the brain do not give rise to any other cell type^53,54^, implying a possible independent functional role for these cells. As such, OPCs are now suggested to be a distinct glial cell type^55^ implicated in brain plasticity through roles such as integration of synaptic activity^56^ and mediation of long term potentiation^57^. Additionally, there is evidence directly implicating the loss of this cell type with emergence of depressive-like behaviour^58^. The data from this study indicate that in MDD OPCs are not only precursor cells for oligodendrocytes, but act as an independently functioning cell type.

Our paper builds on numerous pieces of convergent evidence pointing to the role of synaptic plasticity in the etiopathogenesis of major depressive disorder. In addition, these data expose several other possible paths toward deconvoluting the molecular and cellular changes underlying depression. With time and new insights into similar single nucleus transcriptomic alterations observed in other key brain regions associated with depression, we will be in a better position to forage new avenues for therapeutic interventions.

## Acknowledgements

GT holds a Canada Research Chair (Tier 1) and a NARSAD Distinguished Investigator Award. He is supported by grants from the Canadian Institute of Health Research (CIHR) (FDN148374 and EGM141899), and by the *Fonds de recherche du Québec-Santé* (FRQS) through the Quebec Network on Suicide, Mood Disorders and Related Disorders.

We acknowledge the expert help of the Douglas-Bell Canada Brain Bank staff (Josée Prud’homme, Maâmar Bouchouka and Annie Baccichet), and the technology development team the McGill University and Genome Quebec Innovation Centre (Yu Chang Wang). We would like to thank Liam O’Leary for his artistic inputs in figure creation.

## Author Contributions

CN conceptualized, performed and wrote the manuscript, MM performed experiments, bioinformatics and wrote manuscript, MS and JFT performed bioinformatics and reviewed the manuscript, NM contributed to tissue processing, data interpretation and manuscript preparation, JR provided technical single-cell expertise and experimental support, aided in manuscript preparation, GT provided general oversight, including in experimental design, data interpretation and manuscript preparation.

## Materials and Methods

### Subjects: Postmortem brain samples

This study was approved by the Douglas Hospital Research Ethics Board, and written informed consent from next-of-kin was obtained for each subject. Postmortem brain samples were provided by the Douglas-Bell Canada Brain Bank (www.douglasbrainbank.ca). Frozen grey matter samples were dissected from the left cerebral hemisphere of Brodmann Area 9 (dlPFC). Brains were dissected by trained neuroanatomists and stored at −80 °C. For each individual, the cause of death was determined by the Quebec Coroner’s office, and psychological autopsies were performed by proxy-based interviews, as described previously^1^. Cases met criteria for MDD whereas controls were individuals who died suddenly and did not have evidence of any axis I disorders (Table 1). Post mortem interval (PMI) represents the delay between a subject’s death and collection and processing of the brain.

### Nuclei isolation and capture

50 mg of frozen tissue was dounced in 3 mL of lysis buffer, 10 times with a loose pestle and an additional 5 times with the tight pestle. The sample was left to lyse in a total of 5 mL of buffer for 5 min, after which 5 mL of wash buffer was added and swirled. The sample was passed through a 30 μm cell strainer and spun for 5 min at 500 *g*. This step was repeated for a total of two filtering steps. After pelleting, the nuclei are resuspended in 5-10 mL of wash buffer by pipetting up and down 8-10 times. After 3 washes, the nuclei were resuspended in 1 mL of wash buffer and mixed with 25 % Optiprep™, and layered on a 29 % optiprep cushion and spun for 30 min at 10,000 *g*. Nuclei were resuspended in wash buffer to achieve a concentration of ~1×10^6^ nuclei/mL.

We used the 10x Genomics^®^ Chromium™ controller for single cell gene expression to isolate single nuclei for downstream bulk RNA library preparation. We strictly followed the protocol as outlined by the user guide (CG000052), with the exception of loading concentration, which we increase by 30% as we assessed the capture of nuclei to be slightly less efficient than cell encapsulation. We aimed to capture ~3000 nuclei per sample. This system only allows for a maximum of 8 samples per capture run. As such, we required multiple batches to collect the individual nuclei for all 34 samples (6 batches). Samples 250 and 251 performed poorly, we therefore, carried out the capture on two separate chips and sequenced twice combining the data from both runs for the final analysis.

### Sequence Alignment and UMI Counting

A pre-mRNA transcriptome was built using the cellranger mkref (Cellranger version 2.0.1) command and default parameters starting with the refdata-cellranger-GRCh38-1.2.0 transcriptome and as per the instructions provided on the 10X Genomics website. Reads were demultiplexed by sample index using the cellranger mkfastq command (Cellranger v2.1.0). Fastq files were aligned to the custom transcriptome, cell barcodes were demultiplexed, and UMIs corresponding to genes were counted using the cellranger count command and default parameters.

### Data Transformation for Secondary Analysis

The unfiltered gene barcode matrices for each sample were loaded into R using the Read10X function in the Seurat R package (version 2.2.0, 2.3.0)^2^. Cell names were modified such that the subject name, batch, and biological condition were added to them. Seurat objects were created corresponding to each sample using the CreateSeuratObject function with the imported unfiltered gene-barcode matrices provided as the raw data. Individual Seurat objects for each sample were combined into one object using the MergeSeurat function sequentially. No filtering or normalization was performed up to this step. Since this is a single nucleus dataset, all mitochondrial genes that are transcribed from the mitochondrial genome were removed, along with genes not detected in any cell.

### Barcode and Gene Filtering

Based on the distribution of nGene (total number of genes detected in each cell) for the total dataset (assessed by summary and hist R^3^ functions), barcodes that were associated with less than 110 detected genes were removed. Based on the distribution of nUMI (total numbers of UMIs detected in each cell), the top 0.5 % of barcodes were also excluded as most likely being multiplets rather than single nuclei as there was a very sharp increase of nUMI from 16,393 at the 99.5^th^ percentile to 102,583 at the maximum.

Next, the distribution of nUMI for the remaining barcodes was fit with three normal distributions using the normalmixEM function from the mixtools^4^ package (Supplementary Fig. 1c). The rationale was that, the filtered barcodes contain a population of low quality “noise” barcodes that have a very low nUMI on average, a population of non-neuronal cells that have an intermediate nUMI and a population of neuronal cells that have a high nUMI. Based on the fitting of the normal distributions, only the barcodes with a high probability (> 0.95) of belonging to either the putative “non-neuronal” or putative “neuronal” distributions, and a low probability (<0.05) of belonging to the “noise” distribution were retained for further analysis (Supplementary Fig. 1c-d). 78,886 cells and 30,062 genes were retained.

### Data Processing and Dimensionality Reduction

The UMI counts were normalized to 10,000 counts per cell and converted to log scale (Seurat function NormalizeData). The batch, condition, and subject information was added as meta data to the final Seurat object; nUMI and batch were regressed out using the ScaleData function. The Seurat FindVariableGenes function was used with default selections and cut-offs as follows: x.low.cutoff = 0.003, x.high.cutoff = 2, y.cutoff = 1. This resulted in a list of 2135 highly variable genes, which excludes lowly expressed genes (below 25^th^ percentile), very highly expressed genes, and selects only the top 10 % of genes in terms of the scaled dispersion. These highly variable genes were used to calculate 100 principal components. Based on the PC elbow plot of the standard deviation of the PCs (Supplementary Fig. 2a), the first 50 PCs were retained for use in downstream analysis.

### Clustering by Gene Expression

The FindClusters function was applied with a resolution of 2.5 and identified in 44 initial clusters. The goal of clustering is to sort nuclei by cell type so that all remaining gene expression variation within clusters is not related to cell differentiation processes. Prior to the advent of single nuclei expression profiling, cell types were identified by observing differences in cell morphology, behaviour, and anatomic location. It is fairly straight-forward to sort single nuclei expression profiles into known cell types according to the expression levels of marker genes that differentiate between these cell types. However, it is very unlikely that all cell types have been identified so we must rely on nuclei clustering to uncover as-yet unknown cell types. Unfortunately, the number of clusters obtained from the clustering algorithm is somewhat arbitrary because clustering depends on the settings of several parameters, and there is no consensus on how they should be set. Although clusters obtained using reasonable default settings usually correspond to known biological cell types, some clusters may appear to potentially identify entirely new cell types or splinter existing cell types into multiple subtypes. Deciding if the clusters really do identify new cell types can be difficult or may even be impossible from available data.

To address this issue, we used tools in the Seurat package to sequentially combine any clusters that were not sufficiently distinct from each other. In particular, after performing initial hierarchical clustering of the graph-based clusters (BuildClusterTree), we assessed the nodes of the dendrogram using a random forest classifier (AssessNodes) and then merged together any nodes which were in the bottom 25 % of the dendrogram (using the branching.times function from the ape R package^5^) and had an out-of-bag-error of more than 5 %. We then repeated this clustering and merging process for the nuclei within each terminal node until none of the remaining nodes fulfilled our cut-off criteria (Supplementary Fig. 2b). The resulting set of 30 clusters were then characterized in terms of known markers genes of all major, well-defined brain cell types (Supplementary Fig. 2c-d). For refining identification of excitatory neuron types, we combined and re-clustered a set of excitatory clusters with highly correlated gene expression profiles (R > 0.95) (Supplementary Fig. 3a-c) to get 33 final clusters.

### Cluster Annotation

Genes used as markers for major cell-types and layer-specificity are listed below. Inhibitory neuron subtypes were annotated based on expression of canonical inhibitory interneuron markers SST, PVALB, and VIP where possible. Excitatory neuron subtypes were annotated with some level of layer specificity based on expression of layer specific markers. We also characterised clusters in terms of all genes differentially expressed between clusters (FindAllMarkers function, bimodal test, logfc.threshold of log(2), other parameters set to default) (Supplementary Table 11).

### Major cell-type markers

**Macrophage/ Microglia:** MRC1, TMEM119, CX3CR1; **Endothelial:** CLDN5, VTN; **Astrocytes**: GLUL, SOX9, AQP4, GJA1, NDRG2, GFAP, ALDH1A1, ALDH1L1, VIM; **OPCs:** PTGDS, PDGFRA, PCDH15, OLIG2, OLIG1; **Oligodendrocytes:** PLP1, MAG,MOG, MOBP, MBP; **Excitatory neurons:** SATB2, SLC17A7, SLC17A6; **Inhibitory neurons:** GAD1,GAD2, SLC32A1; **Neurons:** SNAP25,STMN2, RBFOX3.

#### Layer-specific markers

**L2:** GLRA3; **L2-3:** LAMP5, CARTPT; **L2-4:** CUX2, THSD7A; **L2-6:** RASGRF2, PVRL3; **L3-4**: PRSS12; **L4-5**: RORB; **L4-6:** GRIK4; **L5**: KCNK2, SULF2, PCP4, HTR2C, FEZF2: **L5-6**: TOX, ETV1, RPRM, RXFP1, FOXP2; **L6:** SYT6, OPRK1, NR4A2, SYNPR, TLE, NTNG2, ADRA2A

### Purification of Clusters for Differential Expression

While the level of cluster purity we achieved from the above steps was comparable to that of previously published studies, a preliminary assessment of differential expression between our biological conditions within each cell type clusters indicated that we needed further “cluster purification” steps to remove even very small contaminating populations of doublets or misclassified cells. Without such purification uneven presence of contaminating cells can result in false positives in the differentially expressed genes identified. Our purification approach comprised of calculating a median gene expression profile for all our clusters, calculating the correlation of the gene expression of each cell, with the median profile of its cluster (considering only the top 865 genes whose median expression was highly variable, that is had a variance of > 0.25 across the different cluster) and selecting cells with high correlation. This was done by fitting bimodal normal distributions to the total distribution of correlations in the cluster to identify low and high correlation peaks. Cells were retained only if they had a low probability of falling in the low correlation peak (p < 0.25) and a high probability (p > 0.75) of falling in the high correlation peaks (Supplementary Fig. 7).

### Differential Expression Analysis

Differential expression analysis between the cases and controls was performed using linear mixed models implemented in the lme4^6^ and lmerTest^7^ R packages. Mixed models were necessary in order to account for dependencies between nuclei obtained from the same subject. Biological condition and number of UMIs were included in models as fixed effects and the subject and batch as random effects. A false discovery rate (FDR) of 0.1 was used to detect differentially expressed genes within each cell type.

### Pseudotime trajectory using Monocle

For oligodendrocyte developmental trajectory assessment, the data for cells belonging to the five clusters in the oligodendrocyte lineage (Oligos_1, Oligos_2, Oligos_3, OPCs_1, OPCs_2) were used to create a separate Seurat object using the SubsetData function. The most variable genes for these clusters alone were identified using the FindVariableGenes function and the following parameters: x.low.cutoff = 0.003, x.high.cutoff = 3, y.cutoff = 1 (giving a total of 895). The Seurat object was imported into a CDS (CellDataSet) object using the Monocle^8^ function importCDS.

Estimation of size factors and dispersions was performed (using the estimateSizeFactors and estimateDispersions Monocle functions) on the CDS object using default parameters. Dimensionality reduction was then performed using reduceDimension, with reduction_method set to DDRTree. The 895 variable genes identified as above were used for ordering the cells into a trajectory with the orderCells function. The pseudotime trajectory was then plotted with plot_cell_trajectory (Fig. 3a), and the change in expression of genes known to be involved in oligodendrocyte development were plotted using plot_genes_in_pseudotime (Fig. 3b-h).

differentialGeneTest was applied separately to oligodendrocyte lineage cells from control subjects and MDD cases with fullModelFormulaStr = "~sm.ns(Pseudotime)". This allows us to model the expression of each gene as a function of pseudotime. All genes detected in at least one cell in the respective group were compared and their changes across pseudotime were assessed. A q-value cut-off of < 0.01 was used to identify genes associated with pseudotime. The overlapping and non-overlapping genes were identified by comparing the lists obtained for the two groups (Fig. 3a).

### Correlations between differentially expressed genes

The average expression profile per subject within each cluster for each of the 96 differentially expressed genes was calculated using the AverageExpression function in Seurat. Only subjects that contributed cells to all of 16 clusters with differentially expressed genes were retained (13 controls and 6 MDD cases). The correlation coefficient between the expression of every pair of genes was calculated independently for the controls and the MDD cases. One gene (*ZFP36*) with zero average expression in all 6 retained cases was dropped because correlation could not be calculated, leaving 95 genes for further analysis. To compare correlation coefficients between cases and controls, correlation coefficients were transformed to Fisher z-scores using the fisherz function of the R psych^9^ package and a comparison z-score derived using the following formula:

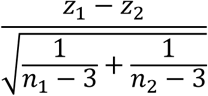

where z_1_ denotes the z-score for the cases, n_1_ the number of cases, z_2_ the z-score for the controls, n_2_ the number of controls. The resulting z-score for the comparison was converted to a two-tailed p-value (Supplementary Fig. 9). P-values were not corrected for multiple testing.

For assessing the overall strength of correlation differences between clusters we used Fisher’s method for combining p-values for each pair of clusters. These combined p-values were used to scale the links in the Circos^10^ plot depicting overall correlation differences (Fig. 4c).

### Cell deconvolution

Expression data from (dbGaP:phs000424.v8.p1)^11^ was used as reference signatures for annotated cell types. UMI counts for each cell were converted to transcripts per million (TPMs) in order to account for the varying sequencing depth of each cell and sample. Average expression levels were calculated for each cell type-specific cluster defined in the paper.

Cluster-specific gene expression profiles were obtained by summing the UMI values of all 24301 genes common to our dataset and the reference for each nucleus in each cluster and converting the sums to TPMs. R package, DeconRNASeq v1.18.0^12^ was used to deconvolute these cluster-specific profiles. Using the data from^11^as reference, we were able to estimate the cell type composition of our clusters.

### Data Availability

Raw sequencing data, annotated gene-barcode matrix, and lists of cells used for differential gene expression analysis are accessible on the MGSS server: http://mgss.cs.mcgill.ca/snRNAseq_paper/

